# Three’s Company: Anthropogenic factors shape co-occurrence in a meso-carnivore community in a semi-arid shared landscape of western India

**DOI:** 10.1101/2024.04.04.588082

**Authors:** Divyajyoti Ganguly, Arjun Srivathsa, Divya Vasudev, Uma Ramakrishnan

**Affiliations:** Wildlife Biology and Conservation Program, National Centre for Biological Sciences, TIFR, Bengaluru, India; National Centre for Biological Sciences, TIFR, Bengaluru, India; Wildlife Conservation Society–India, Bengaluru, India; Conservation Initiatives, Guwahati, India

**Keywords:** free-ranging dogs, interspecific interactions, habitat use, human impacts, carnivores, shared landscapes, tropics

## Abstract

1. Human modification of natural landscapes accelerates species extinctions, promotes biological invasions, coerce range shifts, and restructure biotic communities. Mammalian meso-carnivores, considered sentinels of human-use landscapes, may experience increased mortality through road collisions and interactions with free-ranging dogs. Yet they may also benefit through resource subsidisation and decreased competition with larger carnivores. Our understanding of meso-carnivore community structure in modified ecosystems remain severely limited.
2. We examined a meso-carnivore community comprising golden jackal *Canis aureus*, jungle cat *Felis chaus*, Indian fox *Vulpes bengalensis* and desert cat *Felis lybica ornata* in a semi-arid landscape of Kachchh, western India. First, we mapped their potential habitats and the extent of fragmentation as a measure of vulnerability. We then assessed spatial, temporal and fine-scale spatio-temporal co-occurrence and responses to anthropogenic influences in the meso-carnivore community. We used a combination of camera-trapping and indirect sign surveys within an occupancy modelling framework to generate insights on species space-use and interactions.
3. The most fragmented habitat, open savanna, was positively associated with the occurrence of golden jackal, Indian fox and desert cat. Golden jackal spatially co-occurred with jungle cat and Indian fox with desert cat. Indian fox and desert cat showed spatial avoidance of golden jackal and jungle cat. Meso-carnivores did not show any spatial responses to free-ranging dogs, but showed varied responses to human infrastructure, i.e., settlements, roads and wind turbines, generally co-occurring less with each other in areas with high anthropogenic influence.
4. Spatially co-occurring meso-carnivores also overlapped substantially in the temporal dimension. Golden jackal and jungle cat showed high temporal overlap with free-ranging dogs, while Indian fox and desert cat appeared to avoid them. Temporally overlapping species generally showed fine-scale spatio-temporal aggregation.
5. Our results reveal species-specific responses to anthropogenic factors in the meso-carnivore community, often increasing overlap across space and time in a resource restricted landscape. Future loss of native savanna habitats and an increasing human footprint can elicit competition driven loss of sensitive species leading to homogenisation of the meso-carnivore community. Our findings highlight the importance of considering species-specific responses as well the complex interplay amongst species to conserve multi-carnivore systems across shared, resource-constrained landscapes.

## 1 INTRODUCTION

Human activities have led to the modification of at least 95% of all global ecoregions, primarily driven by land-use change and infrastructure development (Kennedy et al., 2018). These perturbations to natural environments cause the loss and fragmentation of wildlife habitats, leading to changes in forage availability, reproductive success and spatial genetic patterns (Sage, 2019). Further, this can have cascading impacts, such as species extinctions and range shifts, ultimately leading to reorganisation of biotic communities (Tylianakis et al., 2008). The removal or suppression of native species creates vacant ecological niches, making the habitat more susceptible to invasion by better-adapted or competitively superior species (Hui et al., 2016). This can lead to altered competitive dynamics within these communities. Specifically, the removal of a dominant species may allow subordinate species to thrive, forging novel or modified interactions, and an altered community structure and dynamics (Rodriguez, 2006; Tylianakis et al., 2008; Chow-Fraser et al., 2022).

The impacts of human-driven modification can be particularly pronounced in naturally resource-restricted landscapes (e.g., hot and cold deserts) characterised by extreme temperatures, water scarcity and high seasonal fluxes in resource availability (Abere & Oguzor, 2011). There is potential for changes in land-use to mitigate the impact of resource limitation by replacing a fundamentally fluctuating ecosystem with one that is more stable and with an increased resource pool, thereby reducing interspecific competition (Faeth et al., 2005). For instance, avian diversity closer to farms was higher, and reduced with increasing distance to farms in the sand dunes of the Middle East suggesting greater resource access in modified environments, allowing more species to coexist (Khoury & Al-Shamlih, 2006). Conversely, scarce resources in such landscapes may become further restricted, intensifying competitive interactions (Laity, 2009). This might exert additional stress on species, which are already under strong selection pressures to survive harsh environmental conditions (Abere & Oguzor, 2011). Keehn and Feldman (2018) report, for example, that diversity across several taxonomic groups was lower within wind farms, as compared to the outside areas, in the Californian desert ecosystem, indicating higher costs to coexistence near human infrastructure. These altered competitive interactions can manifest in changing community structure, or modified behaviour.

Carnivore communities are typically shaped by competitive forces, and they co-occur with humans across transformed and novel landscapes, globally (see Maddox, 2003; Athreya et al., 2013; Karanth et al., 2017). There are two contrasting hypotheses that may explain species’ responses to human activities in such scenarios. The ‘human super predator hypothesis’ posits that anthropogenic elements, such as humans and human infrastructure (e.g., roads), act as the ‘ultimate predator’ by suppressing carnivore populations through direct or indirect mechanisms (Frid & Dill, 2002; Darimont et al., 2015). In addition, proximity to humans also exposes carnivores to increased competition, predation and disease transfer from free-ranging dogs (*Canis lupus familiaris*) and other companion species (Knobel et al., 2014; Vanak et al., 2014). Alternatively, the ‘human shield hypothesis’ suggests that human proximity confers reduced competition for some species from larger and/or dominant carnivores, coupled with greater access to resources such as human-subsidised foods (e.g., near settlements; Berger, 2007; Bateman & Fleming, 2012). Yet, carnivores may still avoid humans by adapting their activity periods to strictly nocturnal hours (Wang et al., 2015). Such species-level responses can alter the nature of inter-specific interactions. For instance, increased overlap between species in space or time driven by their mutual avoidance of anthropogenic influences may enhance interference competition (Wang et al., 2015; Parsons et al. 2019). These processes can change the structure and composition of multi-carnivore systems as a whole depending on species dominance ranks.

Mammalian meso-carnivores are thought to be better-adapted to anthropogenic landscapes because of their smaller size (<15kg) and area requirements, eclectic dietary habits and higher population growth rates (van Valkenburgh, 2007; Roemer et al., 2009; Marneweck et al., 2022). Meso-carnivores are typically more diverse than large carnivore assemblages, and play complementary ecological roles through predation, scavenging, and seed dispersal (Roemer et al., 2009; Hämäläinen et al., 2017). Given this premise, recent studies have called for their recognition as sentinels for understanding global change impacts on biodiversity across shared landscapes (Marneweck et al., 2022). Within meso-carnivore communities, competitive superiority is typically determined by traits such as body size and sociality (Polis et al., 1989; Palomares & Caro, 1999) – larger and/or more gregarious species suppress smaller and solitary ones, leading to population declines and/or shifts in space use followed by temporal activity or diet profiles of the latter (Newsome & Ripple, 2015; Ferreiro-Arias et al., 2021). When partitioning across these axes is not feasible, species may adapt fine-scale behavioural mechanisms, differing more subtly in the time of their use of the same space (Karanth et al., 2017). In landscapes under anthropogenic influences, in addition to the above-mentioned traits, species’ habitat and dietary niche breadths emerge as important determinants of competitive abilities (Clavel et al., 2011; Monterroso et al., 2020; Suraci et al., 2021) – smaller size along with habitat and diet generality confers a competitive advantage over larger sizes with habitat and diet specialty (Suraci et al., 2021).

Most of our understanding on how meso-carnivores respond to anthropogenic influences comes from studies in temperate regions or locations with relatively low human densities. Tropical countries support high diversities of meso-carnivores, juxtaposed with pronounced impacts of high human population densities on native ecosystems (Di Minin et al. 2016; Marneweck et al. 2021). India ranks among the highest in terms of global carnivore species richness –– 23% of all mammalian carnivores, most of which are meso-carnivores –– alongside the world’s largest human population (>1.4 billion; Srivathsa et al., 2022).

Resource-restricted arid and semi-arid tracts of the country, which still support a high diversity of meso-carnivores, currently face pressures from infrastructure development and rapid land conversions (Madhusudan & Vanak, 2022). Given this background, we sought to understand how a meso-carnivore community responds to human activities in a tropical semi-arid landscape under substantial pressure. Focusing on one such landscape in north-western India, we examined (i) spatial determinants of habitat use by a meso-carnivore community, (ii) the effects of free-ranging dogs on meso-carnivores’ occurrence and co-occurrence patterns, (iii) impacts of human infrastructure (settlements, roads and wind turbines) on the meso-carnivores, and (iv) the potential temporal and fine-scale avoidance that could enable spatially co-occurring species (meso-carnivores and free-ranging dogs) to coexist.

## 2 MATERIALS AND METHODS

### 2.1 STUDY AREA AND SPECIES

The study was conducted across 856 km^2^ of the Kachchh district, Gujarat, India, encompassing parts of the Abdasa, Nakhatrana, and Lakhpat sub-districts (Fig. 1). It is semi-arid landscape characterised by high summer temperatures (45°C) and an average annual rainfall of 412 mm. The habitat is a mosaic of native savannas, invasive *Prosopis juliflora* patches, and croplands. The main crops cultivated in the area include millets, green gram, castor, and groundnut (Anon, 2011). Humans, livestock, and free-ranging dogs (henceforth, dogs) occur across the landscape; herders accompanied by their dogs typically use the savannas to graze livestock during the daytime. In addition to livestock rearing and agriculture, charcoal production from burning *Prosopis juliflora* is a common source of livelihood. Large portions of this landscape are wrongly classified as “wastelands” under Indian land-use policy and consequently receives minimal legal protection or conservation attention (Anon, 2019; Madhusudan and Vanak 2022). Settlements, roads and wind turbines are the major human infrastructures in the region.

**Fig. 1:**
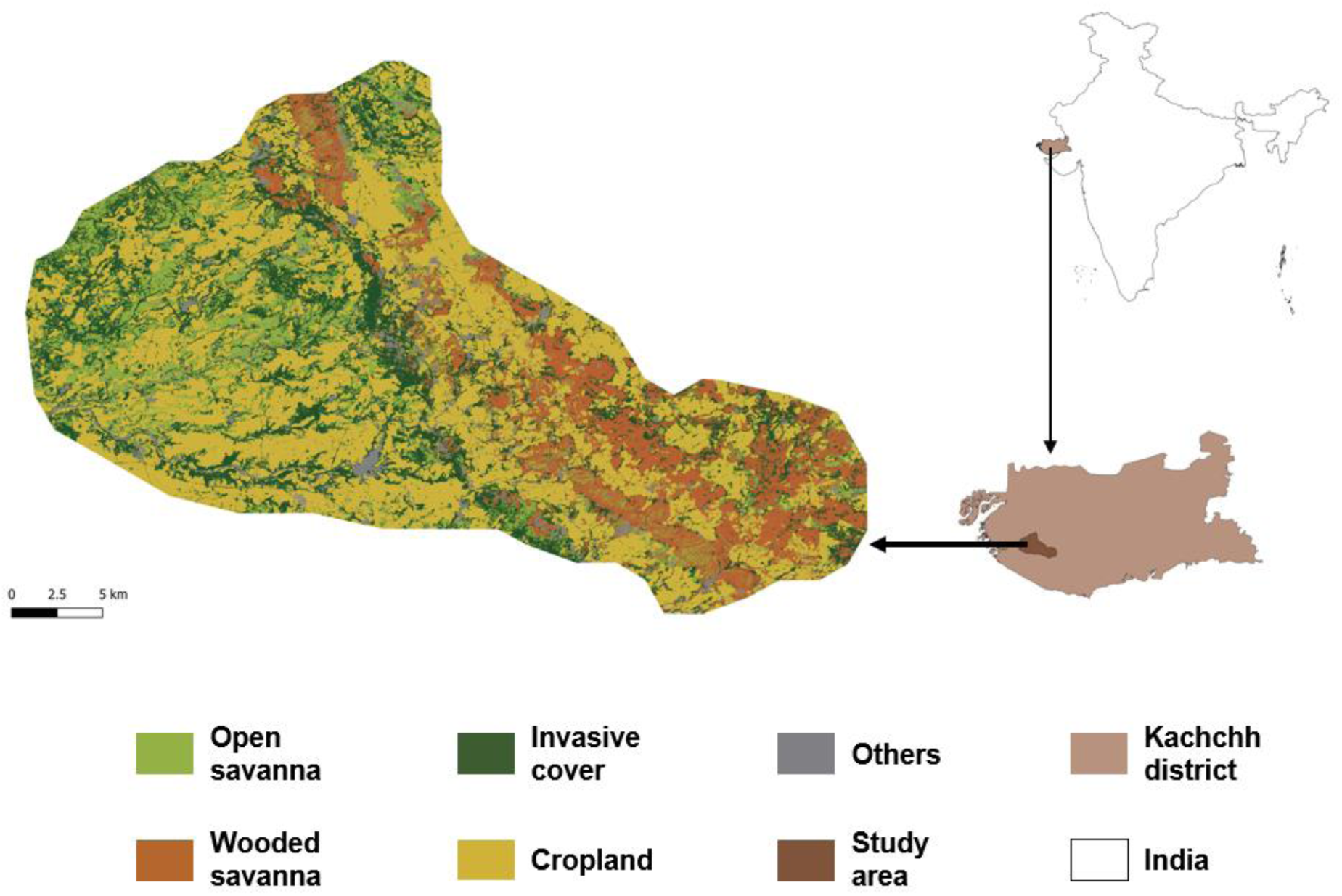
Map of study area in mainland Kachchh – (top right) Location of Kachchh in India; (bottom right) Study area within Kachchh; (left) Land Use Land Cover (LULC) map of the study area generated as part of this study.

The landscape supports a diverse meso-carnivore community with four species of small wild felids, three species of wild canids and one mustelid. Persistent retribution killings led to the local extinction of Indian peninsular grey wolf (*Canis lupus pallipes*), the native large carnivore and apex predator (Vasava, 2015). For this study, we focus on four meso-carnivore species that are relatively common in the landscape: (i) golden jackal (*Canis aureus*), a widely distributed habitat and diet generalist; (ii) jungle cat (*Felis chaus*), a widely distributed habitat generalist and a diet specialist; (iii) Indian fox (*Vulpes bengalensis*), a widely distributed grassland specialist and a diet generalist; and (iv) desert cat (*Felis lybica ornata*), a range-restricted scrubland specialist and also a diet specialist (Castelló, 2018, 2020). These species represent a continuum of traits; they can be ranked in decreasing order of body weight: golden jackal > jungle cat > desert cat > Indian fox; increasing order of habitat specialisation: golden jackal < jungle cat < Indian fox < desert cat; or, increasing order of dietary specialisation: golden jackal < Indian fox < jungle cat and desert cat. Taken together, we expected golden jackals to have a competitive advantage over all species whereas desert cats were likely to be subordinate; we expected jungle cats and Indian foxes to be competitively intermediary.

### 2.2 DATA COLLECTION AND PROCESSING

#### 2.2.1 LAND COVER MAPPING AND CHARACTERISING THE LANDSCAPE

We first generated a land use/land cover map for the study area, given inaccuracies in existing maps. Ground-truthed points were collected using a hand-held GPS device (Garmin eTrex 10) and the Android OS application ‘GPS Essentials’. We supplemented these data with additional points that we visually identified from satellite imagery using Google Earth Pro, and the most recent Bhuvan Land Use Land Cover (LULC) data published in 2019 (https://bhuvan-app1.nrsc.gov.in/thematic/thematic/index.php). We identified the following land-use categories to be the most prominent in our landscape and relevant to our study objectives: native savanna ecosystems including (i) open savanna (grass dominated) and (ii) wooded savanna (native *Acacia sp.* dominated); (iii) invasive cover (*Prosopis juliflora*); and (iv) croplands (Fig. S1). The final training dataset had 105 open savanna points, 69 wooded savanna points, 159 invasive cover points, 200 cropland points and 99 points in the ‘others’ category (water bodies, mines/quarries, solar power plants, factories and settlements). We used a supervised classification framework to generate a land use/land cover map, and used a confusion matrix to assess classification accuracy (Fig. 1; see Supplementary File for additional details).

To assess the status, in terms of availability and fragmentation, of the potential meso-carnivore habitats, we calculated overall proportion of each habitat type, number of patches of each habitat type, mean patch size and level of aggregation of habitat patches using the software FRAGSTATS (McGarigal et al., 1995). The former is an index of habitat availability, while the rest are indicators of configuration, or fragmentation. The level of habitat patch aggregation was measured using the ‘Clumpiness Index’, which ranges from –1 (patch maximally disaggregated) to +1 (patch maximally aggregated).

#### 2.2.2 SAMPLING FRAMEWORK AND FIELD SURVEYS

To detect meso-carnivore presence, we used a combination of camera trap and indirect sign surveys under an occupancy framework (MacKenzie et al., 2002). This study design allowed for maximising spatial coverage and obtaining adequate information on the meso-carnivores, while circumnavigating logistical and time constraints. Sampling units were defined by dividing the study area into a grid network of 4 km^2^ cells (n = 202), considering the average annual home range sizes of three (golden jackal: 12.3 km^2^, jungle cat: 5.9 km^2^ and Indian fox: 4.3 km^2^) of the four study species for which the information was available (Katna et al., 2021). The short survey period (90 days) ensured geographic closure.

Field surveys were conducted during the dry season from January to May 2022. Following a checkerboard format, we ear-marked every alternate cell for camera trapping (n = 101) and carried out the surveys following a block-shifting approach for logistical convenience. One camera trap (Browning/Bushnell/Moultrie) was deployed per cell near the cell centroid, typically at locations with evidence of meso-carnivore presence such as scats/tracks— the maximum deployment distance was 500 m from the centroid. The cameras were active up to 14 days. Every 24-hour interval (12 am to 12 am) was treated as one independent sampling occasion; the total sampling effort therefore ranged up to 14 occasions, or temporal replicates for each grid cell. Details on camera trap data processing are provided in the Supplementary File.

All the cells in the study area were marked for indirect sign surveys (n = 202). Sampling was carried out along the diagonals of cells to record signs of carnivore presence. This field design was chosen because: (i) it ensured adequate spatial coverage of each cell, and (ii) we could reasonably assume that the sample routes were random with respect to animal movement, thus ensuring spatial independence of consecutive sampling segments (Fig. 2).

**Fig. 2:**
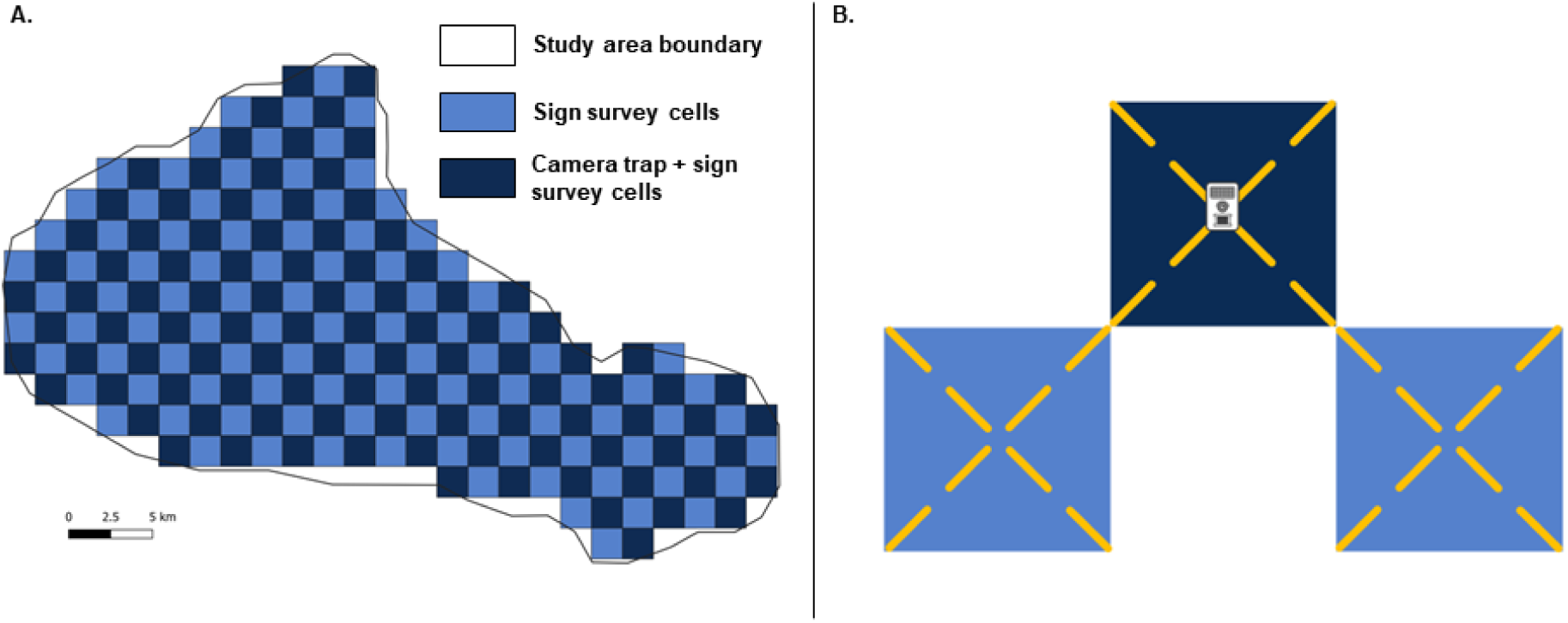
Study area sampling scheme– A. Proposed sampling scheme (202 cells); B. Layout of camera trap stations – one camera trap at the centre of the cell (for every alternate cell), and indirect sign survey routes – one cell with eight segments where each segment is 700 m in length (for all cells).

We considered every 700 m segment along the cell diagonals (one-fourth the length of each diagonal) to comprise one spatial replicate; each cell could therefore have 8 spatial replicates. Track marks of meso-carnivores were difficult to identify in the field. We therefore relied only on genetically confirmed scats for our analysis. Details on field and laboratory procedures for genetic species identification are provided in the Supplementary File.

### 2.3 ANALYTICAL FRAMEWORK

For each species we constructed detection histories (0s for non-detection and 1s for detection in every replicate for each cell), using data generated from camera surveys and the indirect sign surveys. For the camera trap data, we used program *camtrapR* (Niedballa et al., 2016) and for the scat survey data, we manually coded detection histories for golden jackal, jungle cat and Indian fox (desert cat data were excluded as we were unable to unambiguously establish their presence from signs even using genetic tools, see Supplementary File). The two detection histories (one for camera trap and the other for sign survey) thus yielded for each species were appended for further analysis.

Our modelling workflow had three steps, each delving deeper into the spatial responses of meso-carnivore to human impacts (Fig. 3). The first step involved modelling species habitat use with landscape or local habitat variables as predictors. In the second step, we examined spatial co-occurrence patterns among the meso-carnivores, also testing the effect of dogs—a potential competitor in human modified landscapes— on species occurrence and co-occurrence. In the third step, the impacts of human infrastructure on species occurrence and co-occurrence were investigated. We compiled geospatial and ground-based variables (habitat related: land cover, terrain ruggedness; encounters of dogs; and the presence/extent of human infrastructure) as potential covariates that may influence meso-carnivore occurrence and co-occurrence (Table 1 and Supplementary File). All non-categorical covariates were z-transformed and checked for collinearity prior to analysis. For those species without spatial segregation, we compared time activity and within this subset, for those species that showed temporal overlap (Ridout & Linkie, 2009), we analysed fine-scale behavioural interactions using time-to-encounter at camera trap locations (Karanth et al., 2017).

**Fig. 3:**
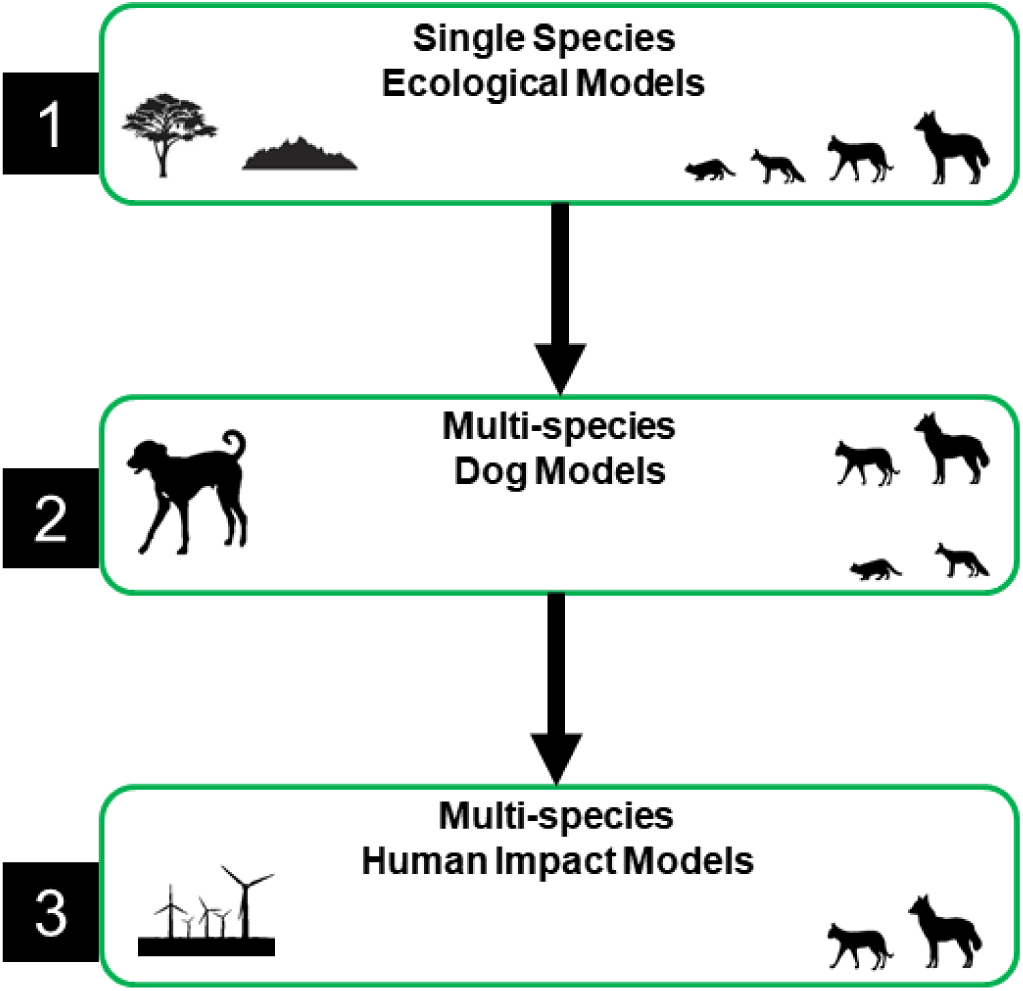
The three-step modelling workflow - first step involved modelling habitat associations (1), second step involved modelling co-occurrence and the effect of dogs on meso-carnivore occurrence and co-occurrence (2) and the third step involved modelling the effects of human impacts as measured through distance to the nearest settlement, density of metalled roads and number of wind turbines on occurrence and co-occurrence (3).

**Table 1:** Expected covariate effects on the focal meso-carnivore species in Kachchh based on current knowledge of their ecology and natural history.

|  |  | Golden jackal | Jungle cat | Indian fox | Desert cat |
| --- | --- | --- | --- | --- | --- |
| Proportion of habitat cover | Open savanna (OS) | 0 | 0 | + | - |
|  | Wooded savanna (WS) | 0 | 0 | + | + |
|  | Invasive cover (IN) | 0 | 0 | - | + |
|  | Cropland (AG) | 0 | 0 | - | - |
| Terrain ruggedness index (TRI) |  | 0 | + | - | - |
| Free-ranging dog encounter rate (DOG) |  | + | - | - | - |
| Distance to nearest settlement (DT) |  | - | - | + | + |
| Density of metalled roads (RD) |  | + | - | - | - |
| Number of wind turbines (WT) |  | + | - | - | - |
| Explanation |  | neutral response to habitat features — habitat generalist, associate positively with dogs — both feed on human derived foods, and with higher human impacts — represent areas with higher scavenging opportunities on wastes/carcasses | neutral response to habitat features — habitat generalist, avoid dogs — potential intraguild predators, and higher human impacts — represent disturbed sites | positive association only with savanna habitats — grassland specialist, avoid dogs — potential intra-guild predators and higher human impacts — represent disturbed sites | positive association with habitats with cover — scrubland specialist, avoid dogs — potential intra-guild predators and higher human impacts — represent disturbed sites. |
| References |  | Castelló, 2018; Devarajan, 2015 | Sunquist & Sunquist, 2017; Castelló, 2020 | Castelló, 2018; Devarajan, 2015; Katna et al., 2021 | Sharma, 1979; Castelló, 2020 |

#### 2.3.2 MESO-CARNIVORE OCCURRENCE AND CO-OCCURRENCE

Before starting our 3–step analysis of meso-carnivore occurrence and co-occurrence, we had to fix the detection parameter. To do this, we used the model described by MacKenzie et al. (2002), which allows for estimating two key parameters: occupancy probability (ψ) and detection probability (*p*). To model detection, we constructed a model set for each species where ψ was modelled as a function of two predetermined habitat covariates and detection either (i) remained constant (null), or (ii) varied as a function of survey method (camera trap or indirect sign survey) — to account for the integration of the two types of survey data (Chaudhary et al., 2020) or (iii) varied as a function of method and associated survey effort. The covariates in the best supported model were used to model detection probability (*p*) across all steps described in the following section.

The first step in our workflow involved modelling species habitat use via single-season single-species occupancy models; MacKenzie et al., 2002). For each species we constructed model sets with combinations of habitat covariates (the proportion of each land cover type per cell) and terrain ruggedness to model ψ. A maximum of two habitat variables were included in any given candidate model so as to avoid multicollinearity. We used the best-supported model based on AICc scores to predict site occupancy (interpreted as probability of habitat use), and make inference on habitat affinities for each species in the landscape (Burnham & Anderson, 2002).

Next, we examined pair-wise co-occurrence using the multi-species occupancy model described by Rota et al. (2016), which allows for parameterising exclusive species presence (marginal occupancy) and species co-occurrence (conditional occupancy) separately. For any two co-occurring species, in addition to occupancy (ψ) and detection probability (*p*), this model allows for estimating a species interaction parameter (*f12*). Here, a negative sign on the estimated interaction term *f12* indicates spatial avoidance, 0 indicates no significant overlap or avoidance, and a positive value indicates spatial overlap. We note that spatial overlap (or lack thereof) in occurrence patterns cannot provide inference on causality in terms of competitive exclusion (Blanchet et al 2020). However, our approach allowed us to explicitly account for imperfect detection of species, and explore patterns of spatial overlap– over and above shared species-habitat relationships.

To test for the impact of dogs on each species and on species interactions, we constructed a set of five candidate models (step 2, Fig. 3) for each of the six species pairs. These were (i) the null model (ψ for both species and *f12* are held constant), (ii) a model where dog presence does not affect occurrence or co-occurrence, but species occurrence is allowed to spatially vary based on our inferences on species-habitat relationships from Step 1 (ψ for both species modelled as a function of habitat covariates; *f12* is held constant), (iii) where dog affects only species occurrence, but not their interactions (ψ for both species affected by dogs and habitat covariates; *f12* is held constant), (iv) dog affects only species co-occurrence, but not their occurrence (ψ for both species affected by habitat covariates; *f12* is affected by dogs) and (v) dog affects species occurrence as well as co-occurrence (ψ for both species affected by dogs and habitat covariates; *f12* is affected by dogs). We retained the covariate combination(s) from the best-supported model(s) based on AICc scores from this step for subsequent analyses.

In step 3 (Fig. 3), we constructed 23 models for each species pair with all possible additive combinations of settlements, roads and wind turbines as potential covariates affecting meso-carnivore occurrence (after accounting for species-habitat associations) and co-occurrence for all species-pairs. The final inferences were based on the models with the best relative fit (ΔAICc <2; Burnham & Anderson, 2002) in each modelling step. All the occupancy analyses were performed in program R using package *unmarked* (Fiske & Chandler, 2011; R Core Team, 2022).

#### 2.3.3 TEMPORAL OVERLAPS

Competing species which are spatially segregated need not avoid each other in time – temporal avoidance facilitates coexistence between spatially co-occurring species. Hence, temporal activity patterns and overlap were assessed between meso-carnivore pairs and between meso-carnivores and dogs, only for those pairs with no evidence for spatial avoidance (based on the multi-species occupancy analyses). For this, species activity kernels were generated based on the time-stamp of photo-detections from camera traps. The extent of temporal overlap was measured as 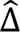, or the overlap in the area under the kernel curves in the activity of two species in each pair, calculated using R packages *camtrapR* and *overlap* (Ridout & Linkie, 2009, Niedballa et al., 2022). The metric 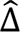, ranges from 0 to 1, with values closer to 0 indicating lower overlap in activity patterns, i.e., higher temporal partitioning. We interpret values with upper bound in SE <=0.5 to imply low overlap and values with lower bound in SE >0.5 as high overlap. We calculated the modified 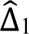 for species pairs with small sample sizes (<50 records); for other species pairs, we calculated and interpret 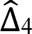, which is regarded as a better alternative for larger sample sizes (Ridout & Linkie, 2009).

#### 2.3.4 FINE-SCALE SPATIO-TEMPORAL INTERACTIONS

Species pairs that showed spatial overlaps at the 4-km^2^ grid-cell level and temporal overlaps in terms of 24-hour diel activities could still be avoiding each other at finer scales. To investigate this, we assessed fine-scale spatio-temporal interactions between those species pairs that showed evidence of temporal association in the previous step. We adapted the multi-response permutation procedure followed by Karanth et al. (2017), which compares observed minimum time-to-encounter in camera traps between species pairs, with an expected minimum time-to-encounter under an assumption of randomness (i.e., neither aggregation nor avoidance). We estimated random time-to-encounter by shuffling photo-detections, while maintaining for each species, the total number of detections and diel activity patterns. To account for the block-shifting method followed in our study, wherein all camera traps were not active across our entire sampling period, we randomised time of encounter, and then randomly selected a date on which the camera trap was active. We then calculated minimum time-to-encounter. We repeated this process 1000 times for each species pair, and generated a distribution of median expected minimum times-to-encounter. For each species pair, if the observed median time-to-encounter was less than the expected (random) value, we interpreted it as species aggregation; a value higher than that expected through randomisation implies species avoidance (Karanth et al, 2017).

## 3 RESULTS

Field surveys were carried out in 163 grid cells, of which we carried out camera trap surveys in 95 cells, and sign surveys in 68 grid-cells. Of these, 56 cells were sampled using both methods (Fig. S3). The surveys entailed a total effort of 950 camera-trap days and 374 km of indirect sign surveys. We generated 464 independent carnivore photo-encounters and genetically screened 127 carnivore scat samples, which resulted in the detection of 12 wild carnivore species (see Table S3), free-ranging cats (*Felis catus*) and dogs.

### 3.1 STATUS OF MESO-CARNIVORE HABITATS

Open savannas had the least spatial coverage (∼14%) in the study area. Savanna habitats were highly patchy, and the patches themselves were extremely small and relatively less aggregated in space (Table 2), indicating high fragmentation. Croplands, on the other hand, had the highest proportional coverage (∼44%), and were unfragmented with individual patches being larger and more contiguous (Table 2).

**Table 2:** Class descriptors of the habitats in the study area landscape as computed on FRAGSTATS. The ‘Clumpiness Index’ values range from -1 to 1 where the closer the value to 1, the more aggregated the habitat patches.

| Habitat type | Percentage landscape coverage | Number of patches | Mean patch area (ha) | Clumpiness Index |
| --- | --- | --- | --- | --- |
| Open savanna | 13.63 | 32118 | 0.34 | 0.78 |
| Wooded savanna | 15.37 | 17299 | 0.70 | 0.87 |
| Invasive cover | 22.55 | 33450 | 0.53 | 0.81 |
| Cropland | 43.81 | 6755 | 5.13 | 0.94 |

### 3.2 MESO-CARNIVORE–HABITAT ASSOCIATIONS

The naïve occupancy estimates for golden jackal, jungle cat, Indian fox and desert cat were 0.34, 0.37, 0.15 and 0.25 respectively and the predicted occupancy probabilities (inferred as habitat use) from the invariant model in each case were 0.58 (0.06 SE), 0.54 (0.06 SE), 0.25 (0.05 SE) and 0.47 (0.12 SE) respectively (Fig. 4). The associated detection probabilities predicted from the best supported detection model (Table S4) were 0.19 (0.02 SE), 0.12 (0.02 SE), 0.17 (0.03 SE), 0.06 (0.02 SE) from camera trap surveys; detectability from scat surveys were 0.04 (0.01 SE), 0.16 (0.03 SE) and 0.04 (0.02 SE) for golden jackal, jungle cat and Indian fox, respectively. Golden jackal, Indian fox and desert cat showed a positive association with open savanna habitat (Table S6). Golden jackal and jungle cat showed a negative association with wooded savanna habitat and rugged terrain. Jungle cat showed a weak positive response to croplands (Table S6).

**Fig. 4:**
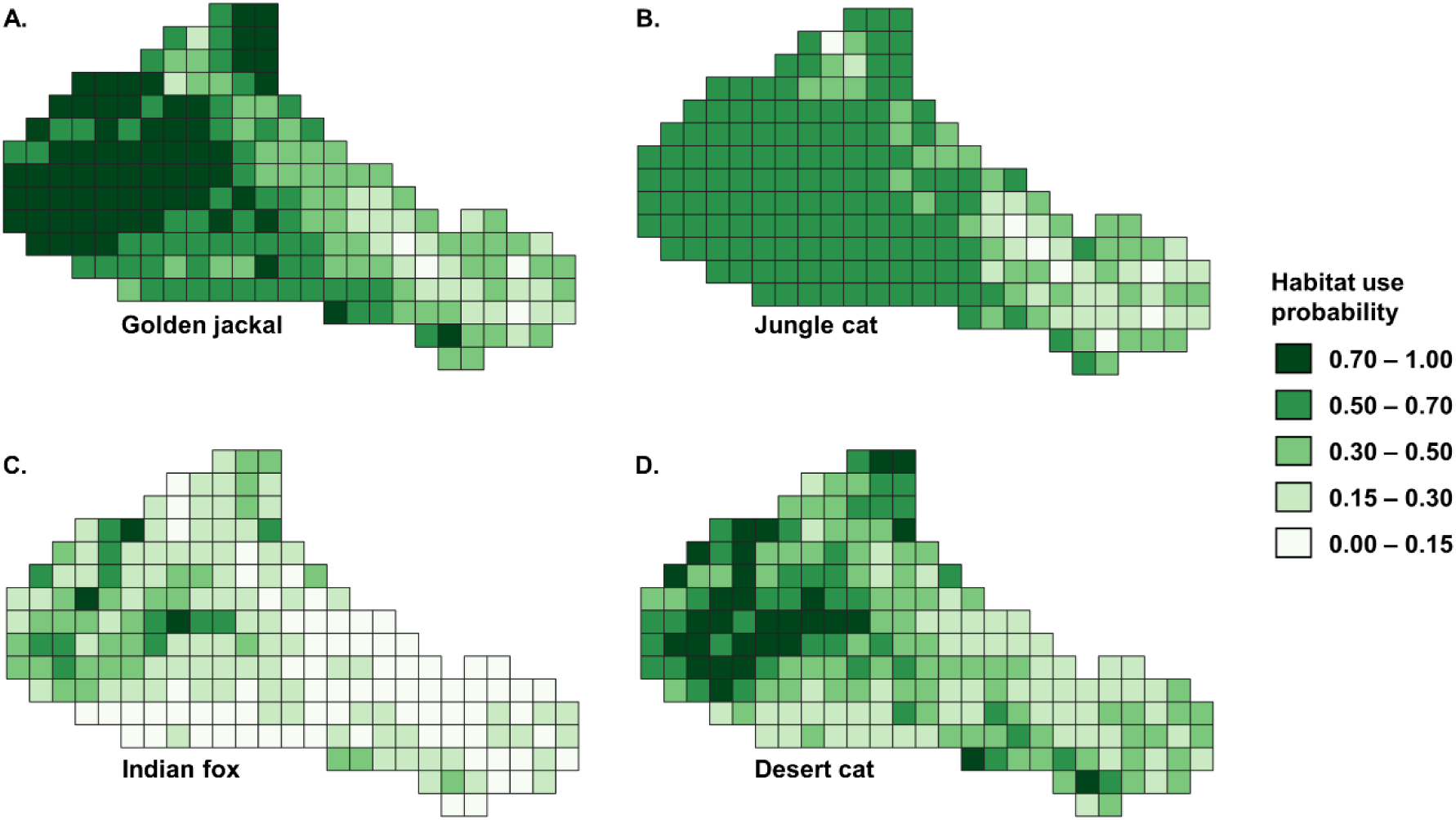
Map of predicted site occupancy (inferred as habitat use) based on the best supported models for the focal meso-carnivores – A. golden jackal (range = 0.12–0.97), B. jungle cat (range = 0.09–0.67), C. Indian fox (range = 0.11–0.75) and D. desert cat (range = 0.23–0.95).

### 3.3 PAIRWISE SPATIAL CO-OCCURRENCE AND IMPACT OF DOGS

Spatial avoidance was observed in four species pairs: golden jackal–Indian fox, jungle cat– Indian fox, golden jackal–desert cat and jungle cat–desert cat (Table S8). Two species pairs, golden jackal–jungle cat (*f12* = 1.17, 0.61 SE) and Indian fox–desert cat (*f12* = 2.16, 1.83 SE) showed positive spatial association. We expected dogs to have a negative effect on all species except golden jackals, for which we expected a positive association (Table 1). None of the models with dogs as a predictor of meso-carnivore occurrence had adequate statistical support (ΔAICc >2; Table 3). Further, dog encounter rates did not show significant effects on any of the pairwise species’ co-occurrence.

**Table 3:** Models with statistical support for the multi-species modelling step to examine the influence of dogs on meso-carnivore occurrence and co-occurrence (AICc < 2). K: number of parameters; AICc: Akaike’s Information Criterion corrected for small sample size; ΔAICc: model AICc–minimum AICc; ModelLik: model likelihood; AICcWt: model weight; LL: log likelihood; Cum.Wt: cumulative model weight. All the covariates were z-transformed prior to analysis. For full list of models, see Table S6.

| Model | K | $AICc$ | $\Delta AICc$ | Model Lik | $AICcWt$ | LL | Cum. Wt |
| --- | --- | --- | --- | --- | --- | --- | --- |
| **Golden jackal (sp1) and Jungle cat (sp2)** |  |  |  |  |  |  |  |
| $\psi_1(OS+WS)$,<br>$\psi_2(WS)$,<br>$f_{12}(\cdot)$ | 10 | 1614.3 | 0 | 1 | 0.57 | -796.42 | 0.57 |
| **Golden jackal (sp1) and Indian fox (sp2)** |  |  |  |  |  |  |  |
| $\psi_1(OS+WS)$,<br>$\psi_2(OS)$,<br>$f_{12}(\cdot)$ | 10 | 1185.65 | 0 | 1 | 0.61 | -582.1 | 0.61 |
| **Golden jackal (sp1) and Desert cat (sp2)** |  |  |  |  |  |  |  |
| $\psi_1(OS+WS)$,<br>$\psi_2(OS)$,<br>$f_{12}(\cdot)$ | 8 | 982.04 | 0 | 1 | 0.61 | -482.18 | 0.61 |
| $\psi_1(OS+WS)$,<br>$\psi_2(OS)$,<br>$f_{12}(DOG)$ | 9 | 983.97 | 1.93 | 0.38 | 0.23 | -481.93 | 0.85 |
| **Jungle cat (sp1) and Indian fox (sp2)** |  |  |  |  |  |  |  |
| $\psi_1(WS)$,<br>$\psi_2(OS)$,<br>$f_{12}(\cdot)$ | 9 | 1170.93 | 0 | 1 | 0.66 | -575.88 | 0.66 |
| **Jungle cat (sp1) and Desert cat (sp2)** |  |  |  |  |  |  |  |
| $\psi_1(WS)$,<br>$\psi_2(OS)$,<br>$f_{12}(\cdot)$ | 7 | 784.93 | 0 | 1 | 0.75 | -384.82 | 0.75 |
| **Indian fox (sp1) and Desert cat (sp2)** |  |  |  |  |  |  |  |
| $\psi_1(OS)$,<br>$\psi_2(OS)$,<br>$f_{12}(\cdot)$ | 7 | 607.46 | 0 | 1 | 0.65 | -296.08 | 0.65 |
| $\psi_1(OS)$,<br>$\psi_2(OS)$,<br>$f_{12}(DOG)$ | 8 | 609.4 | 1.95 | 0.38 | 0.24 | -295.86 | 0.89 |

OS: proportion of open savanna; WS: proportion of wooded savanna; DOG: free-ranging dog encounter rate; $\psi_1$ : occupancy probability of species 1 (sp1); $\psi_2$ : occupancy probability of species 2 (sp2); $f_{12}$ : species interaction parameter; Detection probability was modelled as CT + SS throughout except for pairs with desert cat, where it was kept constant. CT: camera trap; SS: sign survey.

### 3.4 SPATIAL CO-OCCURRENCE AND HUMAN IMPACTS

Impacts of human infrastructure on meso-carnivore occurrence and co-occurrence featured among the top-ranked models (Table 4). As predicted, increasing distance to settlements negatively affected golden jackals and positively affected Indian foxes (Fig. 5A; Table S10). Contrary to expectation, jungle cats showed higher use away from human settlements, while desert cats had higher use closer to settlements. Species pairs were more likely to co-occur away from settlements. The density of metalled roads affected the occurrence of three focal meso-carnivores negatively, except for jungle cat where it was a positive effect (Fig. 5B; Table S10). Co-occurrence between species pairs reduced in four of the six cases when the density of metalled roads increased. The number of wind turbines did not show any discernible impact on the occurrence of canids, whereas the felids seemed to be positively affected (Fig. 5C; Table S10). Higher number of wind turbines reduced spatial association between golden jackal–jungle cat, while increasing chances of co-occurrence between jungle cat–desert cat (*f12* = 2.16, 0.91 SE).

**Fig. 5:**
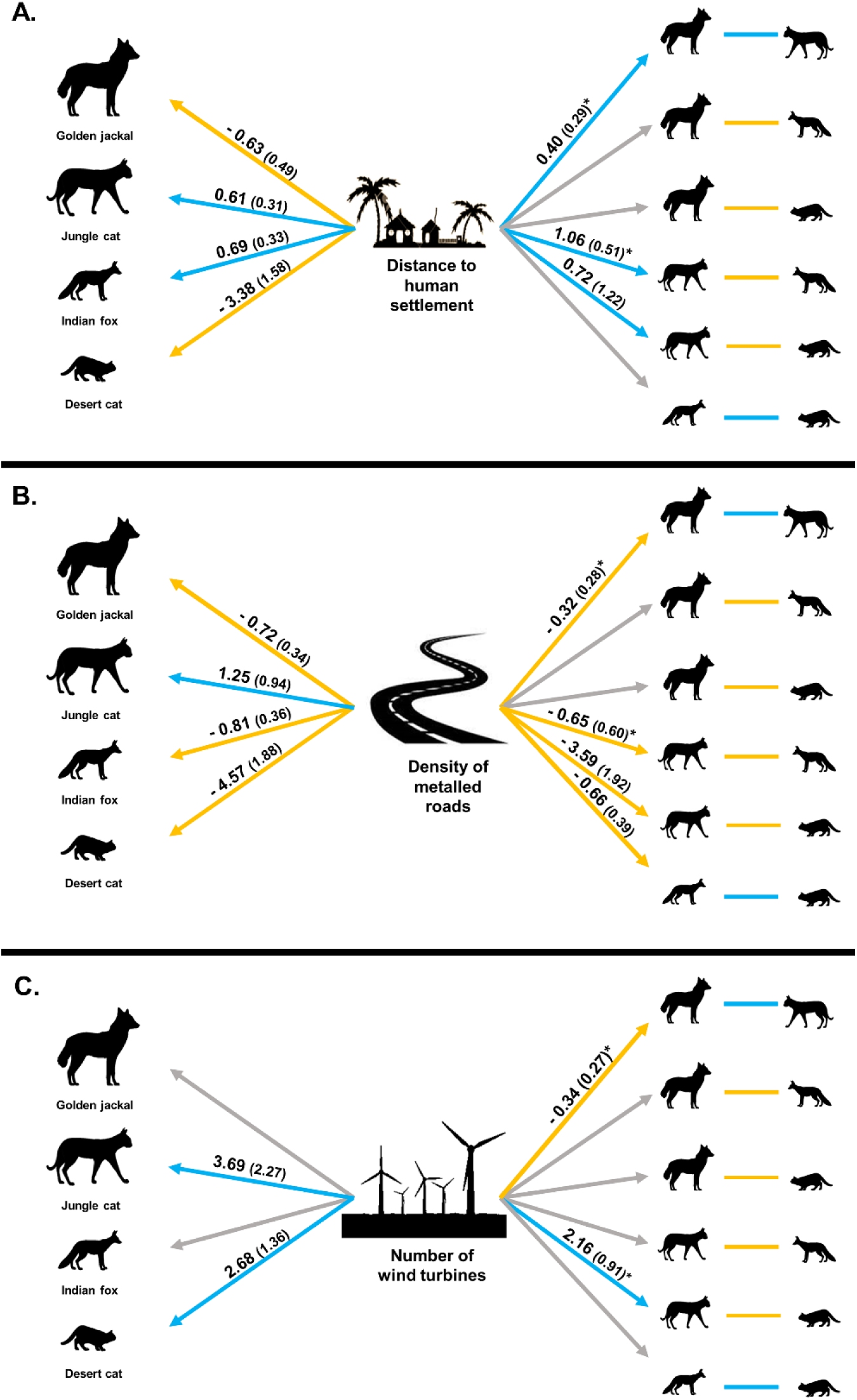
Schematic of meso-carnivore occurrence and co-occurrence as affected by human impacts – A. Settlements, B. Roads and C. Wind turbines. Grey arrows indicate lack of statistical support (AIC > 2 or SE overlapping 0), blue arrows indicate positive association, yellow arrows indicate negative association. Values marked with ‘*’ were taken from models with some statistical support and where it first appeared (ΔAICc < 2) but not from the best supported model (Table S9).

**Table 4:** Models with statistical support for the multi-species modelling step to examine the influence of human infrastructure on meso-carnivore occurrence and co-occurrence (AICc < 2). K: number of parameters; AICc: Akaike’s Information Criterion corrected for small sample size; ΔAICc: model AICc–minimum AICc; ModelLik: model likelihood; AICcWt: model weight; LL: log likelihood; Cum.Wt: cumulative model weight. All the covariates were z-transformed prior to analysis. For full list of models, see Table S9.

| Model | K | $AICc$ | $\Delta AICc$ | Model Lik | $AICc$ Wt | LL | Cum. Wt |
| --- | --- | --- | --- | --- | --- | --- | --- |
| **Golden jackal (sp1) and Jungle cat (sp2)** |  |  |  |  |  |  |  |
| $\psi_1(OS+WS+RD)$,<br>$\psi_2(WS+RD)$, $f_{12}(.)$ | 12 | 1613.53 | 0 | 1 | 0.17 | -793.73 | 0.17 |
| $\psi_1(OS+WS)$, $\psi_2(WS)$,<br>$f_{12}(.)$ | 10 | 1614.3 | 0.76 | 0.68 | 0.11 | -796.42 | 0.28 |
| $\psi_1(OS+WS)$, $\psi_2(WS)$,<br>$f_{12}(DT)$ | 11 | 1614.65 | 1.11 | 0.57 | 0.1 | -795.45 | 0.38 |
| $\psi_1(OS+WS)$, $\psi_2(WS)$,<br>$f_{12}(WT)$ | 11 | 1615 | 1.47 | 0.48 | 0.08 | -795.63 | 0.46 |
| $\psi_1(OS+WS)$, $\psi_2(WS)$,<br>$f_{12}(RD)$ | 11 | 1615.16 | 1.63 | 0.44 | 0.07 | -795.71 | 0.53 |
| $\psi_1(OS+WS)$, $\psi_2(WS)$,<br>$f_{12}(WT+DT)$ | 12 | 1615.25 | 1.71 | 0.42 | 0.07 | -794.58 | 0.6 |
| $\psi_1(OS+WS+DT+RD)$,<br>$\psi_2(WS+DT+RD)$, $f_{12}(.)$ | 14 | 1615.32 | 1.79 | 0.41 | 0.07 | -792.24 | 0.67 |
| **Golden jackal (sp1) and Indian fox (sp2)** |  |  |  |  |  |  |  |
| $\psi_1(OS+WS+RD)$,<br>$\psi_2(OS+RD)$, $f_{12}(.)$ | 12 | 1180.68 | 0 | 1 | 0.42 | -577.3 | 0.42 |
| **Golden jackal (sp1) and Desert cat (sp2)** |  |  |  |  |  |  |  |
| $\psi_1(OS+WS+WT+RD+DT)$,<br>$\psi_2(OS+WT+RD+DT)$, $f_{12}(.)$ | 14 | 975.83 | 0 | 1 | 0.52 | -471.29 | 0.52 |
| **Jungle cat (sp1) and Indian fox (sp2)** |  |  |  |  |  |  |  |
| $\psi_1(WS+DT)$, $\psi_2(OS+DT)$,<br>$f_{12}(.)$ | 11 | 1167.8 | 0 | 1 | 0.2 | -572.02 | 0.2 |
| $\psi_1(WS)$, $\psi_2(OS)$, $f_{12}(DT)$ | 10 | 1168.27 | 0.48 | 0.79 | 0.16 | -573.41 | 0.36 |
| $\psi_1(WS)$, $\psi_2(OS)$, $f_{12}(RD+DT)$ | 11 | 1169.05 | 1.25 | 0.53 | 0.11 | -572.65 | 0.46 |
| $\psi_1(WS)$, $\psi_2(OS)$, $f_{12}(WT+DT)$ | 11 | 1169.69 | 1.89 | 0.39 | 0.08 | -572.97 | 0.54 |
| **Jungle cat (sp1) and Desert cat (sp2)** |  |  |  |  |  |  |  |
| $\psi_1(WS+WT+RD+DT)$,<br>$\psi_2(OS+WT+RD+DT)$,<br>$f_{12}(WT+RD+DT)$ | 16 | 781.03 | 0 | 1 | 0.2 | -371.03 | 0.2 |
| $\psi_1(WS)$, $\psi_2(OS)$, $f_{12}(WT+RD)$ | 9 | 781.87 | 0.83 | 0.66 | 0.13 | -380.87 | 0.33 |
| $\psi_1(WS+WT+DT)$,<br>$\psi_2(OS+WT+DT)$, $f_{12}(WT+DT)$ | 13 | 782.4 | 1.36 | 0.51 | 0.1 | -375.95 | 0.43 |
| $\psi_1(WS+WT+RD)$,<br>$\psi_2(OS+WT+RD)$, $f_{12}(WT+RD)$ | 13 | 782.46 | 1.43 | 0.49 | 0.1 | -375.98 | 0.53 |
| $\psi_1(WS)$, $\psi_2(OS)$, $f_{12}(WT+RD+DT)$ | 10 | 782.48 | 1.45 | 0.48 | 0.1 | -379.93 | 0.62 |
| $\psi_1(WS)$, $\psi_2(OS)$, $f_{12}(WT+DT)$ | 9 | 782.71 | 1.67 | 0.43 | 0.09 | -381.29 | 0.71 |
| **Indian fox (sp1) and Desert cat (sp2)** |  |  |  |  |  |  |  |
| $\psi_1(OS)$, $\psi_2(OS)$, $f_{12}(RD)$ | 8 | 606.22 | 0 | 1 | 0.23 | -294.27 | 0.23 |
| $\psi_1(\text{OS}+\text{RD}), \psi_2(\text{OS}+\text{RD}), f_{12}(\cdot)$ | 9 | 607.3 | 1.08 | 0.58 | 0.14 | -293.59 | 0.37 |
| $\psi_1(\text{OS}), \psi_2(\text{OS}), f_{12}(\cdot)$ | 7 | 607.46 | 1.24 | 0.54 | 0.13 | -296.08 | 0.49 |
| $\psi_1(\text{OS}), \psi_2(\text{OS}), f_{12}(\text{WT}+\text{RD})$ | 9 | 608.17 | 1.95 | 0.38 | 0.09 | -294.02 | 0.58 |

OS: proportion of open savanna; WS: proportion of wooded savanna; DT: distance to nearest settlement; RD: density of metalised roads; WT: number of wind turbines; $\psi_1$ : occupancy probability of species 1 (sp1); $\psi_2$ : occupancy probability of species 2 (sp2); $f_{12}$ : species interaction parameter; Detection probability was modelled as CT + SS throughout except for pairs with desert cat, where it was kept constant. CT: camera trap; SS: sign survey.

### 3.5 OVERLAP IN THE TEMPORAL DIMENSION

All four focal species showed nocturnal to crepuscular peaks in activity. Temporal overlaps were assessed between golden jackal–jungle cat and Indian fox–desert cat because they showed spatial aggregation. The temporal overlap between these species pairs was high (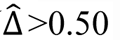; Fig. S4). We also assessed temporal overlap between meso-carnivores and dogs because there was no evidence of spatial avoidance. Golden jackals and jungle cats showed temporal overlap with dogs (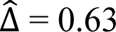, 0.04 SE and 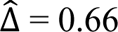, 0.04 SE respectively; Fig. S5); Indian fox and desert cat showed low overlap, implying potential temporal avoidance of dogs (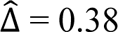, 0.05 SE and 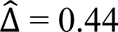, 0.06 SE respectively; Fig. S5).

### 3.6 FINE-SCALE AVOIDANCE AND OVERLAP

We assessed fine scale spatio-temporal interactions (minimum time-to-encounter at camera trap stations) only between those pairs of species for which there was evidence for temporal overlap 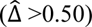. Meso-carnivore species pairs generally showed spatio-temporal aggregation as did dog–golden jackal at finer scales with significantly lower observed time-to-encounters, as compared to random (Fig. 6). Jungle cats avoided dogs with the observed time-to-encounter being significantly higher than expected by chance.

**Fig. 6:**
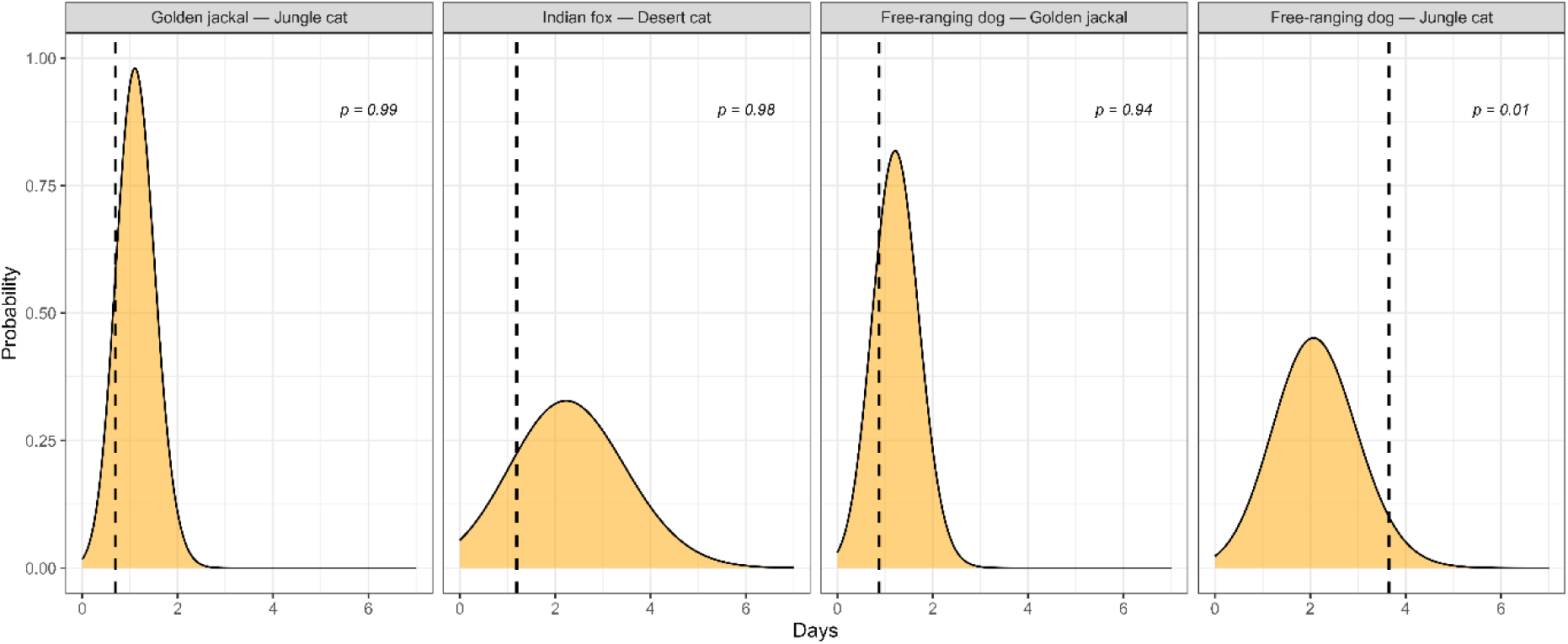
Fine scale interactions, as shown by minimum times-to-encounter among temporally overlapping meso-carnivore and dog pairs. The vertical dashed lines represent median minimum observed time-to-encounter between two species, while the shaded yellow area show randomly simulated times-to-encounters. The *p*-values were calculated as the proportion of randomly generated (expected) times-to-encounter values that are greater than the observed time-to-encounter.

## 4 DISCUSSION

Through a dry season snapshot assessment, we examined spatial ecology and spatio-temporal overlap of species in a meso-carnivore community in a semi-arid, shared landscape in India. We found that open savanna (the most fragmented land cover in the area) was positively associated with the occurrence of three of the four focal species. Interspecific interactions between species pairs varied; four pairs appeared to show some level of avoidance, while the others showed different degrees of aggregation in space, time and at fine spatio-temporal scales. Importantly, species overlap was modified by anthropogenic impacts, particularly the density of metalled roads, impeding species co-occurrence.

### Competitive dynamics among the focal meso-carnivores

Coexistence between carnivores is typically precluded by competition over resources, which results in spatial or temporal or dietary or fine-scale behavioural segregation (Kitchen et al., 1999; Karanth et al., 2017). This may be accentuated in resource-restricted environments where access to food and refuge is limited and unpredictable (Abere & Oguzor, 2011). For instance, use of water sources by kit foxes (*Vulpes macrotis*) was found to be constrained by coyotes (*Canis latrans*) in the arid Mojave Desert, United States (Hall et al., 2021). Instances of competition-driven intraguild mortality peaks when one species is four times larger than the other, and both species are within the same Family (Prugh & Sivy, 2020). We found some evidence for spatial avoidance following a size-based hierarchy where co-occurrence between the smaller Indian fox and desert cat and the larger golden jackal and jungle cat was low.

However, we also found evidence for spatial aggregation between golden jackal–jungle cat and Indian fox–desert cat, where the species pairs have similar body sizes and habitat affinities. These pairs also showed high temporal overlaps and aggregation at fine spatio-temporal resolution (times-to-encounter at individual camera trap locations). Such patterns may be indicative of the clumped nature of common resources, which we were not able to measure or model. It may also be indicative of facilitative interactions whereby meso-carnivore species may enable each other to track unpredictable and ephemeral resources, such as animal carcasses or human-derived foods.

### Interactions between meso-carnivores and dogs

Meso-carnivore suppression through competition with or direct predation by large carnivores is well documented in scientific literature (Linnell & Strand, 2008). Because meso-carnivores tend to alter their spatial and/or temporal activity in response to dogs (almost ubiquitously the world over) researchers have recognised the latter as ‘novel’ apex predators (see Ritchie et al., 2014; Gálvez et al., 2021). Our objective was to test whether dogs show suppression of meso-carnivores akin to large carnivores. We did not find evidence for spatial avoidance between the focal meso-carnivores and dogs; patterns of meso-carnivore co-occurrence were also unaffected. It is possible that spatial partitioning between meso-carnivores and dogs was restricted owing to near-ubiquitous presence of the latter across our landscape. However, avoidance was evident in the temporal dimension, with Indian foxes and desert cats showing low overlap with dogs. These species may also be more vulnerable to mortality because they are smaller than dogs (Devarajan, 2015; Carricondo-Sanchez et al., 2019). Golden jackals and jungle cats, on the other hand, showed relatively higher temporal overlap with dogs. Avoidance between jungle cats and dogs was only apparent at the finest scale, a behavioural response that possibly helps reduce agonistic interactions between the two species.

Considering the human-aided rise in dog populations across South Asia, and that most population control methods have had limited success (Belsare & Vanak 2020; Bhalla et al., 2021), increased interactions between the two might be an impending threat for the meso-carnivores in Kachchh and elsewhere across the world.

### Infrastructure impacts on meso-carnivore ecology and interactions

Human impacts on carnivores manifest as a complex interplay of threats and benefits, thus making it challenging to assess their community-wide effects. Meso-carnivores are drawn to human-inhabited areas due to food subsidies (Ramesh & Downs, 2014; Srivathsa et al., 2020); this proximity often results in increased mortality and reduced fitness (Prugh et al., 2023). Roads, while providing scavenging opportunities from vehicle-caused mortalities, also heighten collision risks to the carnivores themselves (Cook & Blumstein, 2013; Planillo et al., 2018). Over time, this can prompt scavenging carnivores to recognize these locations as ecological traps and avoid areas with substantial vehicular movement (Basille et al., 2013).

We found that golden jackals, Indian foxes, and desert cats avoided areas with higher road densities. Such patterns are not only indicative of reduction in habitat suitability for wild carnivores, but in the long term, can impede connectivity and gene flow (Thatte et al., 2019), underscoring the importance of implementing mitigation measures such as overpasses and underpasses in fragile habitats.

Human settlements lure carnivores not only through food subsidies but also by offering protection from dominant predators (Bateman & Fleming, 2012; Moll et al., 2018). Our focal species showed different tendencies – golden jackals and desert cats appeared to favour sites closer to settlements, while jungle cats and Indian foxes avoided them. This proximity can, for instance, increase interactions between desert and free-ranging/domestic cats, increasing hybridization risks (Gerngross et al., 2022)– another poorly understood aspect of human– carnivore interactions.

Wind turbines cause bird and bat mortalities through blade collisions; these locations thereby attract scavenging carnivores (Lovich & Ennen, 2013). Besides scavenging opportunities, areas around wind turbines can also increase hunting success because of (1) higher prey densities, and (2) lower predator detection caused by turbine noise (Rabin et al., 2006; Thaker et al., 2018). In our study, the felids showed higher use in areas with greater number of turbines. Recent work suggests that meso-carnivores in wind farms can experience physiological stress (Agnew et al., 2016), linked to fitness and population declines (Crespi et al., 2012). Further research on these physiological responses would add to provide an overall picture on the impacts of turbines on meso-carnivores.

Overall, human infrastructure reduced co-occurrence between meso-carnivore species. As road densities increased, co-occurrence decreased for four of six species pairs, one pair showed reduced co-occurrence in areas with more wind turbines, and three pairs were less likely to co-occur closer to human settlements. Human infrastructure sometimes increased co-occurrence between species that usually avoided each other (e.g., jungle cat–desert cat). Considered together, the negative effects of proximity to anthropogenic attributes seemed to outweigh any potential benefits for the meso-carnivores, manifesting either as mutual avoidance or coerced overlap (likely leading to increased competition).

### Vulnerable habitats as refuges in a landscape mosaic

Meso-carnivores are often deemed ecologically ‘plastic’ owing to their widespread distribution (Marneweck et al., 2021). This notion has perhaps contributed to many species being classified as ‘Least Concern’ under the IUCN Red List (Srivathsa et al., 2022). While this status may hold true at a global scale, we show that widely distributed species can still be locally restricted to certain habitats; the occurrence of three of the four focal meso-carnivores was strongly linked to vanishing, fragmented and largely unprotected open savanna habitats. The spatial distribution and configuration of savanna habitat, occurring as a mosaic of grasslands with scattered *Euphorbia sp., Prosopis juliflora* bushes and shrubs, offers meso-carnivores greater access to refuge and prey. For example, chambers formed by *Euphorbia sp.* are used by the desert cat for shelter and denning (Biont, 2022). A bulk of the meso-carnivores’ diet, e.g., golden jackal (∼75%), jungle cat (∼73%), Indian fox (∼44%), is composed of rodents (Sharma, 1979; Mukherjee et al., 2004; Home & Jhala, 2009); grassland mosaics are known to harbour a high diversity and abundance of such rodent prey (see Jayadevan et al., 2018).

In Kachchh, land cover modifications and infrastructure development together indicate an increasing human footprint. Conversion of savanna habitats into croplands is apparent in our analysis (Fig. S6), and a continued growth in the renewable power sector across western and north-western states of India is also anticipated (Sharma, 2011; Dawn et al., 2019). Rural road networks continue to expand with a 69% increase between 2000 and 2015 (https://data.gov.in/). Our results suggest an already changing meso-carnivore community with reduced species co-occurrence near high human impact areas; away from such sites the community is characterised by pronounced inter-specific interactions. For spatially co-occurring species with similar habitat associations, further reductions in nocturnal activity periods to facilitate temporal partitioning or fine-scale behavioural avoidance could entail higher bio-energetic costs in a resource-restricted landscape – coercing high interspecific overlaps. This may promote interference competition among meso-carnivores and between meso-carnivores and dogs. In the long term, as suggested by Seveque et al. (2020), there can be a breakdown of natural community dynamics by filtering out competitively inferior species, impeding species coexistence and simplifying/homogenising the meso-carnivore community. Our study from a densely human-populated tropical habitat adds to previous assessments from elsewhere around the world, the importance of considering species-specific responses *as well as* the complex interactions amongst species in multi-carnivore systems while designing conservation interventions in shared landscapes.

## 5 ACKNOWLEDGEMENTS

We thank the Gujarat Forest Department and Gujarat Biodiversity Board for providing research permits. We thank S. Mahmad, I. Bhatti and B. Thakker for their assistance with field logistics. A. Luhar, A. Jat, R. Jat, M. Jat, J. Juneja, I. Juneja, A. Juneja, A. Mahmad and A. Jat assisted with field surveys; A. Halder and A.K. Pathak helped with data collection; P. Saravanan and A. Sureshbabu helped with data processing.

We are grateful to A. Kashyap and P. Koulgi for their help with the analysis and A. Vishwanathan, J. Ratnam, K. Suryawanshi, J.D. Nichols, J.E. Hines and C. Rota for their inputs at various stages of our study. NCBS–TIFR, Nature Conservation Foundation, and Prakriti Research Fellowship (enabled by CARPE, Grind Master, AITG, Mayuri Kerr, Kedar Shah and Caring Friends Foundation) provided institutional and/or funding support to D.G.

